# Siibra: A software tool suite for realizing a Multilevel Human Brain Atlas from complex data resources

**DOI:** 10.1101/2025.05.20.655042

**Authors:** Timo Dickscheid, Xiaoyun Gui, Ahmet Simsek, Christian Schiffer, Jean-Francois Mangin, Yann Leprince, Viktor Jirsa, Jan G. Bjaalie, Trygve B. Leergaard, Sebastian Bludau, Katrin Amunts

## Abstract

Computational technology opens new possibilities towards understanding the complexity of the human brain, but it requires integrating measurements from different modalities and scales in anatomical context and exposing them in interoperable, actionable form. Especially with growing big data resources, accessing information from different scales and modalities coherently for visual exploration, reproducible analysis and application development remains challenging. We present *siibra*, a tool suite that connects diverse data from cloud resources to reference atlases and coordinate spaces. It supports different use cases by making contents accessible through a web viewer, Python library and HTTP API. Using *siibra* we implemented a Multilevel Human Brain Atlas linking macro-anatomical concepts and their inter-subject variability with measurements of the microstructural composition and intrinsic variance of brain regions, building on cytoarchitecture as a reference and supporting MRI-based and microscopic templates. The atlas is integrated with the EBRAINS research infrastructure. All software and content are openly accessible.

## Main

Decoding the human brain requires consideration of multiple levels of structural and functional organization, using measurements obtained at different spatio-temporal scales with complementary methods. To describe the composition and intrinsic variance of brain regions in conjunction with brain-wide neural network connectivity and inter-subject variability, macro-anatomical concepts need to be linked with microstructural features at cellular resolution. The integration of such comprehensive information requires reference atlases providing consistent and accurate definitions of anatomical structures across a range of spatial resolutions and modalities in multiple reference coordinate systems, and a consistent model for diverse representations of locations in the brain—e.g., brain areas, coordinates, or bounding boxes—that reflect how measurements are linked to anatomy. Such integration of reference atlases with data sets across scales and modalities constitutes a *multilevel atlas*. Its implementation builds on existing concepts, such as those introduced in [55], and imposes new conceptual and technical difficulties. In particular, it must accommodate rapidly evolving computational technologies and automated workflows to ensure efficient collection and processing of the comprehensive information it offers. Although originating from very different data sources, elements of the multilevel atlas must be exposed in well harmonized, interoperable and actionable formats to make them available in a computationally scalable fashion.

Ample data resources exist that describe different aspects of brain organization [53], and a broad range of reference atlases are available reflecting complementary principles of brain organization, typically provided in the form of MRI-scale image volumes or labelled surface meshes at spatial resolutions in the millimeter range derived from in vivo imaging. Integration of information into neuroscience workflows has so far been mostly addressed by casting high-resolution measurements into data representations for macroscopic analyses [51, 16], such as interpolated and averaged feature distributions in surface and voxel formats. Such derivative data make high-resolution features accessible and computable but may reduce the rich regional variance and spatial specificity encoded in the original measurements. In fact, the neuroscience community is increasingly motivated to share primary data with full level of detail and enable their reuse [43]. This is supported by the increasing attention of publishers to this topic, leading to a growing abundance of large primary data resources on different online repositories. However, accessing and using data resources from different scales and modalities at their full level of detail remains a technical challenge. An important reason for this is the lack of coherent spatial and semantic integration. The structural relationships between e.g., microscopic measurements from histological studies and whole-brain neuroimaging measurements are often unclear, and the data may refer to different versions and representations of information in reference atlases. In addition, links between measurements and brain areas, such as the label indices in a parcellation map, get easily lost during analysis workflows. Moreover, the consistent interpretation of spatial locations in the brain requires accounting for localization accuracy, different spatial resolutions, and effects of transformations between reference spaces. Another difficulty is the fundamental difference in handling big data, such as whole brain sections at full resolution and 3D volumes from microscopic studies in the Giga- to Terabyte range, as compared to data being shared in the form of small files.

Progress in obtaining and analyzing large data sets and the need to link the different modalities across spatial scales imply a paradigm shift from downloading files to interacting with cloud resources and services. Researchers are confronted with new and variable data representations and substantially different tool chains. Therefore, to enhance usability and facilitate data use, we need software solutions that expose data from large cloud resources together with classical file-oriented representations in a unified and interoperable fashion and provide operational principles to connect them with workflows for analysis and modeling.

Here we present *siibra* (“<u>S</u>oftware Interfaces for Interacting with <u>BR</u>ain <u>A</u>tlases”), a tool suite designed to tackle these challenges by offering an informatics framework that integrates data from different modalities and resources with reference atlases spanning macroscopic and microscopic resolutions. The tool suite includes an interactive 3D web viewer (*siibra-explorer*), an HTTP API (*siibra-api*), and a Python library (*siibra-python*). The tool suite enables to specify and process various forms of spatial locations in the brain, such as centers of gravity of measured neuroimaging signals, coordinates of functional activations and bounding boxes of regions of interest in brain tissue, and assign them to brain areas defined in reference atlases through multilayered semantic and spatial relationships. This enables anatomical characterization and multimodal profiling of various forms of locations and regions of interest in the brain. Relationships between data, spatial locations and brain areas are modeled as observations with a quantifiable degree of uncertainty and evaluated based on properties such as overlap and correlation of spatial locations and semantic associations according to accepted taxonomies of brain region hierarchies.

We have developed *siibra* to implement a multilevel atlas of the human brain that is integrated with the EBRAINS research infrastructure (https://ebrains.eu) and provides interfaces to additional data repositories. This multilevel atlas combines a complementary selection of existing reference atlases and coordinate spaces with a broad range of multimodal datasets, supporting voxel and surface-based workflows for the whole brain while offering access to underlying data at microscopic detail. The reference spaces include at the macroscopic scale the ICBM 2009c nonlinear asymmetric multi-subject and the Colin27 single-subject average templates [30] as well as surface subspaces (fsaverage and fsaverage6) from the Freesurfer software suite [20, 28]. These are combined with the BigBrain space at microstructural level [4] which represents a 3D reconstruction of 7404 histological sections from an individual postmortem human brain with an isotropic voxel resolution of 20 *×* 20 *×* 20*µm*. A central reference of the Multilevel Human Brain Atlas is the Julich-Brain cytoarchitectonic atlas [5], which includes probabilistic maps of currently almost 300 cortical and subcortical areas; it captures the regional variance in microstructure based on studies in ten postmortem brains. Many of these areas are further annotated as detailed 3D maps across the full stack of BigBrain histological sections [63] using the same methodology in combination with a deep learning workflow [63]. This way, an unambiguous correspondence is established between the in vivo imaging and the ex vivo microscopic scale, which is independent of spatial relationships established between the reference coordinate systems and therefore not affected by the inevitable inaccuracies of image registrations. The cerebral cortex in the BigBrain is further segmented by maps of six isocortical layers based on an automatic classification of 3D cortical intensity profiles [68].

The cytoarchitectonic maps are aligned with maps of superficial as well as long white matter bundles [32, 47] obtained by clustering streamlines from diffusion weighted imaging of a large number of subjects, as well as maps of functional modules at different granularities computed from millions of fMRI scans to provide an efficient reduction and representation of BOLD signals [19]. The data resources connected to the multilevel atlas include detailed measures of cellular (e.g. cell densities; staining profiles) and molecular (e.g. receptor densities; gene expressions) architecture, parcellation-based structural and functional connectivity from neuroimaging cohorts, and an extensive collection of high-resolution histological measurements.

The *siibra* tool suite facilitates the development of this multilevel atlas by enabling efficient access to all content for download, interactive exploration and computational processing—despite high spatial resolutions and a large number of underlying files. It allows tagging all elements of reference atlases and multimodal data features with comprehensive metadata and expose them in interoperable formats to serve a wide range of use cases, such as hypothesis-driven exploration, data analysis, and modeling. The software architecture of *siibra* makes future integration of additional reference atlases, coordinate systems and data resources into the framework straightforward.

We here present the tool suite and demonstrate its functionalities using our implementation of the Multilevel Human Brain Atlas as a challenging use case. By including the underlying executable code or persistent URLs with each figure, the present paper also serves as a practical demonstration of how *siibra* enables fully reproducible workflows that are easily accessible to all researchers.

## Results

In the following, we demonstrate how key functionalities of *siibra* enable the implementation of the Multilevel Human Brain Atlas, linking reference atlases and comprehensive datasets from cloud resources and making the underlying data accessible and actionable for neuroscience applications as illustrated in Figure 1. All data and functionality are openly accessible. References to corresponding executable codes and online resources are provided throughout the text to ensure reproducibility. The work relies on several key concepts, whose distinctions are crucial for understanding the presented results and methodology:

1. We distinguish **individual reference atlases** from the overarching concept of the **Multilevel Human Brain Atlas**. The latter combines complementary reference atlases reflecting different principles of brain organisation, such as the Julich-Brain atlas [5, cytoarchitecture], the deep white matter bundle atlas [32, fibre architecture], or the atlas of dictionaries of functional modes [19, functional architecture] (Figure 1B).
2. Following the AtOM ontology model [42], we assume that each individual reference atlas provides a **terminology** of brain area definitions, often in hierarchical form. Brain areas are delineated in images or surfaces of different **reference coordinate systems**, resulting in **annotation sets**. Annotations may be given in the form of **labelled or statistical maps**, reflecting discrete or continuous annotations respectively. Discrete annotation sets covering the whole brain are often further denoted as **parcellation maps**.
3. The *siibra* tool suite does not store any actual data itself. Instead, all atlas elements and data features point to published data resources, referred to as **siibra content** (Figure 1A). We distinguish **foundational content** linked to *siibra* through explicit definitions collected in *siibra* **configurations**, and **dynamic content** obtained by *siibra* at runtime using **live queries**, i.e., direct software interfaces to cloud resources.

**Figure 1.**
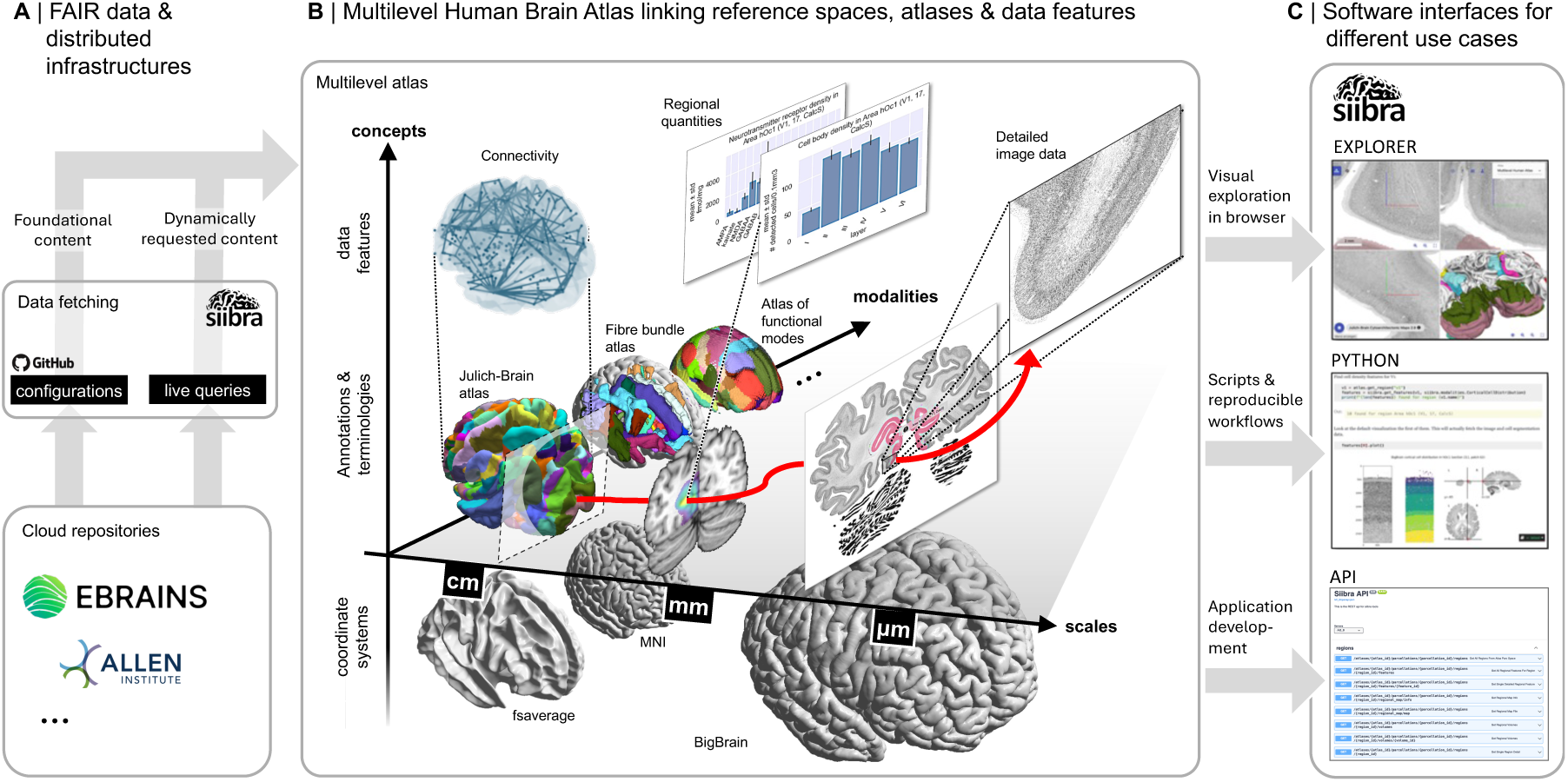
The *siibra* tool suite is used to implement a Multilevel Human Brain Atlas integrating cloud-based reference atlases and datasets from different scales and modalities into an easy-to-use informatics framework for interactive and computational workflows. A: Contents of the multilevel atlas—i.e., reference annotations, terminologies, templates and data features— are curated and stored in cloud repositories, with EBRAINS chosen as the main platform following the openMINDS metadata framework. We distinguish *foundational content*, predefined as versioned collections of configuration files, from *dynamic content* obtained at runtime using automatic “live queries” to APIs of selected cloud repositories. B: The tool suite links comprehensive data resources with reference atlases into the Multilevel Human Brain Atlas. This covers different reference coordinate systems and atlas reference annotations capturing statistical aspects of inter-subject variability as well as microscopic details of individual brains. Brain areas and spatial locations are linked with multimodal datasets to collect measurements of brain organization at different levels. As examples, we illustrate microscopic images capturing brain-wide cell distributions, connectivity matrices encoding functional or structural connections between brain areas, and regional quantities such as tables of neurotransmitter receptor densities. The red arrow illustrates a typical access path from a brain area to a detailed measurement in a relevant dataset, which we cover in detail in the results section. C: The *siibra* tool suite is a modular environment consisting of cloud services and installable software components. It provides access to all contents of the Multilevel Human Brain Atlas in harmonized, actionable data formats through interactive and programmatic user interfaces to serve a broad range of neuroscience use cases.

### Visually guided exploration from full brain networks to cells

The interactive atlas viewer *siibra*-explorer allows visually guided exploration of brain organisation from the macroscopic scale of the brain down to individual cell bodies, and can be accessed with any modern web browser, including mobile touchscreen devices. It integrates very large and detailed image resources (see Supplementary Information, Table S3) into a 3D visualization of the whole brain. In addition, it embeds cross-sectional volumetric and surface views into an interactive workflow that allows to navigate reference atlases in different coordinate systems, and to access multimodal datasets anchored to brain areas or coordinate-based locations. Cross-sectional views can be adjusted in real-time to display arbitrary oblique cutting planes and zoom levels, which is crucial for e.g. studying cortical laminar structure of a highly folded cortical surface. Individual planar views can be maximized to obtain a large field of view, and support layering to co-display brain region maps with underlying image data.

The first example concerns Betz cells in layer V of the primary motor cortex [5, area 4p], illustrated in Figure 2. The workflow starts by selecting the MNI Colin27 template and Julich-Brain as reference atlas, displayed as a cross-planar view combined with a rotatable 3D surface visualization in the corresponding reference coordinate system (Figure 2A). By maximizing one of the cutting planes, a more focused view is obtained and can be adjusted to display an oblique sectioning angle (Figure 2B). When switching to the BigBrain template, precomputed nonlinear spatial transformations are used to preserve location and zoom level while changing the reference coordinate system (Figure 2C), further described in the “Methods” section.

**Figure 2.**
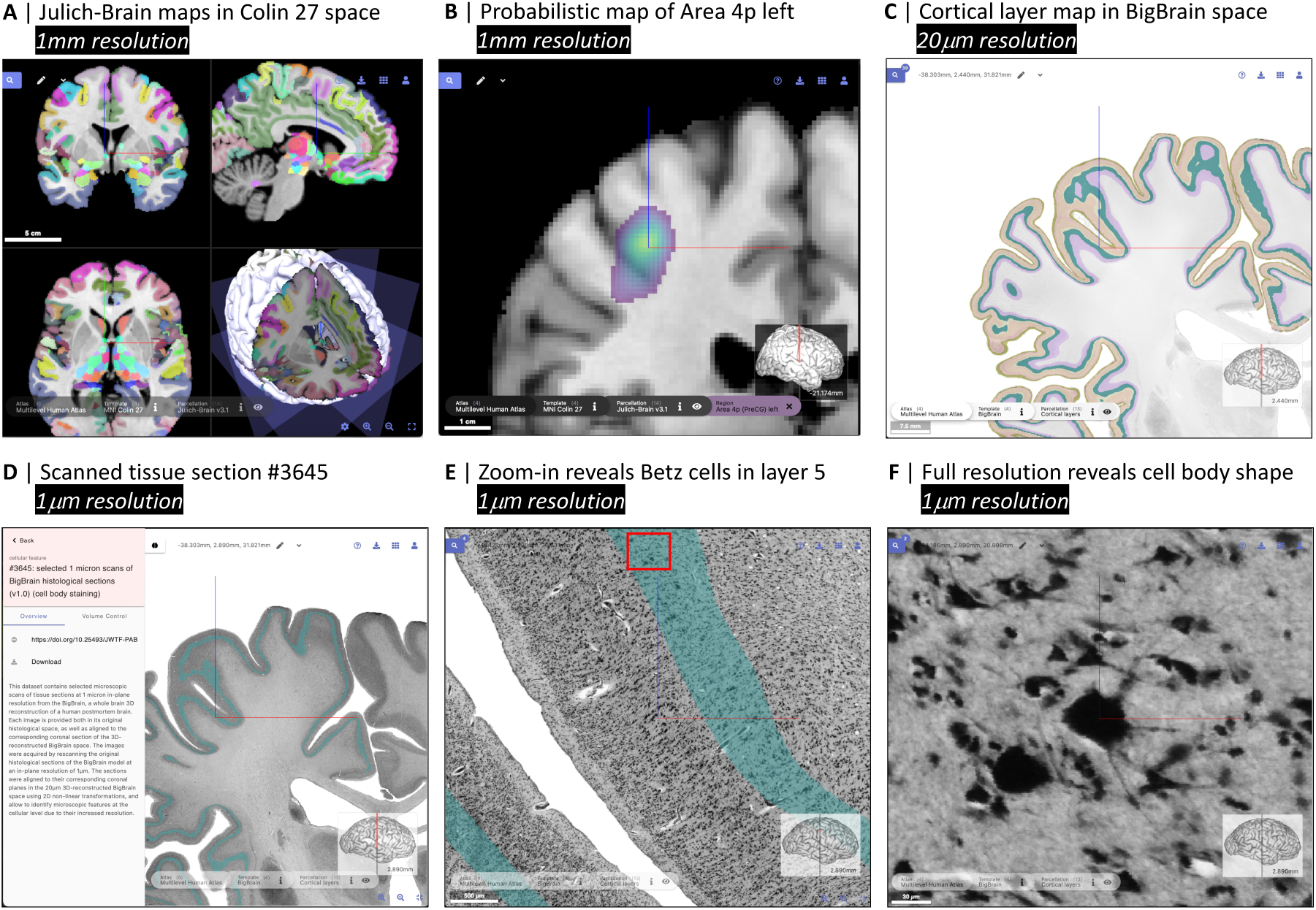
Visually guided exploration of brain structure from the whole brain down to individual cells with *siibra-explorer*. **A, B:** Selection of area 4p in the left hemisphere from the Julich-Brain cytoarchitectonic maps [5] in MNI Colin 27 space [30] brings up its probability map which captures the variability observed in ten postmortem brains. **C:** When switching to the microscopic BigBrain model as a reference space [4], the level of detail is significantly increased while *siibra-explorer* preserves zoom and location. Additional parcellations and anchored image data, including 1*µm* scans of BigBrain sections, are revealed. The parcellation is changed to cortical layer maps [68]. **D:** A 1*µm* scan of a coronal whole brain section [62] is found in the proximity and superimposed with the 3D model. **E, F:** Zooming to full resolution reveals presence and shape of individual cell bodies of Betz Giant cells, clearly located in layer V. Although here only illustrated for D, basic metadata and reference links to publications and online resources are provided for all involved reference templates, atlases and data features. *All steps can be reproduced using the following URLs: A, B, C, D, E, F*.

This mechanism can be used to identify macroscopic structures in MNI space as a starting point for exploring detailed maps of cytoarchitectonic regions and cortical layers in the BigBrain [68], and finding relevant microscopic image data at cellular resolution anchored to this coordinate space (Figure 2D). The viewer allows to select microscopic images and superimpose them with the underlying template, so that they can be explored at full spatial resolution while keeping linked with the whole brain context (Figure 2E,F). The multi-layered views can be reused by bookmarking the browser URL or downloading the underlying zoomedin data, using the download button on the top right.

### Anatomical characterization and multimodal profiling for regions of interest

The *siibra* tool suite enables the assignment of user-defined regions of interest to brain areas from reference atlases and facilitates the retrieval of relevant multimodal measurements. This can be used to further characterize these regions in terms of structure, function and connectivity. In the simplest case, locations are specified as 3D coordinates in a supported reference space (Figure 3A) using *siibra*’s interactive or programmatic location definition options. Associated brain regions are then assigned to brain areas from a reference atlas (Figure 3E) utilizing the available 3D reference maps—e.g. functional, cyto- or fiber architectonic maps—and distinguishing between incidence, correlation and overlap of structures. In the case of statistical maps, where each brain area is typically represented by a separate image volume to account for the overlap in the maps, this might imply to retrieve, load and process several hundred image volumes. To reduce resulting download times and memory requirements, *siibra* uses a sparse data representation for statistical maps which drastically reduces the memory footprint, detailed in the “Methods” section. Associated brain areas can then be used to run data queries and find multimodal data features connected to these structures as shown in Figure 3F. During this process, *siibra* automatically resolves spatial relationships between different reference spaces and, if necessary, warps coordinates using precomputed diffeomorphic transformations.

**Figure 3.**
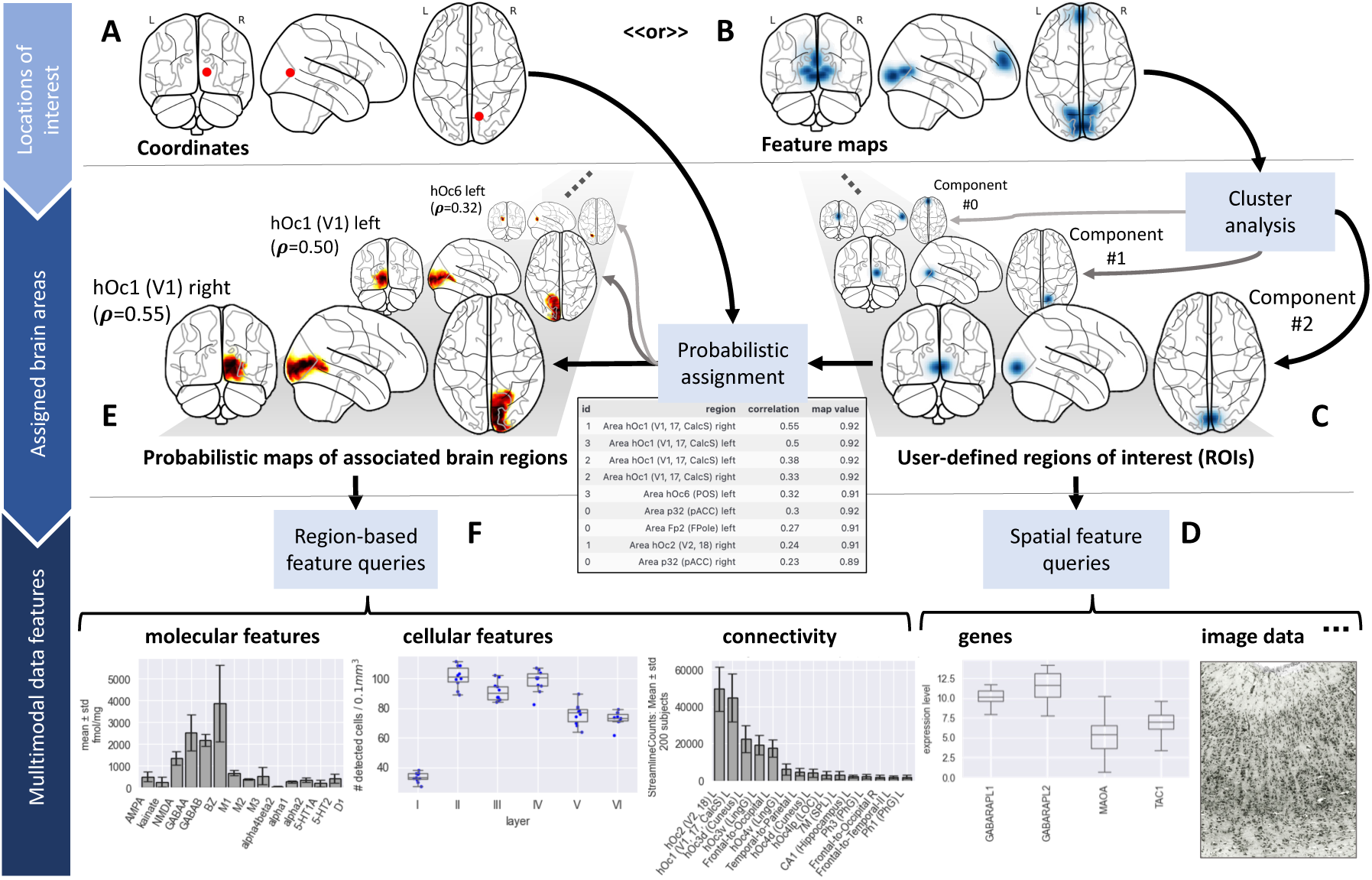
Anatomical characterization and multimodal profiling of regions of interest. *siibra* accepts user-defined location specifications in the form of reference space coordinates with optional certainty quantifiers (A) or feature maps from imaging experiments (B). Feature maps are typically separated into cluster components to obtain clearly localized regions of interest (ROIs) (C). Any region of interest in image form can be used to run spatial feature queries and extract multimodal data features co-localized with ROIs (D). In addition, locations of interest are assigned to brain areas using probabilistic maps from functional, cyto- or fiber architectonic reference atlases (E), distinguishing incidence, correlation and overlap of structures. The resulting associated brain areas reveal additional multimodal data features characterizing the ROIs (F). *The Figure can be reproduced using the notebook available at* https://siibra-python.readthedocs.io/en/v1/examples/tutorials/2025-paper-fig3.html.

The current set of data features in the Multilevel Human Brain Atlas are grouped into the categories cellular, molecular, fibres, functional, connectivity and macrostructure. An overview is provided in the Supplementary Information, Table S3. Their content includes combinations of image, tabular and numerical array data. Tabular data structures include cortical density measures from histological labelings, “fingerprints” encoding means and standard deviations of densities from tissue samples, structural and functional connectivity matrices, or external data such as gene expression levels from the Allen human brain microarray data [33, 1]. Connectivity matrices are parcellation-grouped numbers and lengths of streamlines obtained from tractography of diffusion MRI data as well as correlations of resting-state and task-specific functional time series extracted from different neuroimaging cohorts. Image features include tissue sections, volumes of interest from high-resolution imaging experiments, and in vivo neuroimaging scans. Histological image data are typically anchored to the BigBrain template, since it provides the necessary level of detail to embed microscopy data. Links between data features and atlas elements are further discussed in the “Methods” section.

In addition to coordinates, a region of interest may be specified by an image volume (Figure 3 B), e.g., an activation map from a functional neuroimaging study. By processing segments or certain intensity ranges, images can be directly used for spatial feature queries (Figure 3 C) to access spatially anchored datasets such as co-registered 2D and 3D image data, or regional quantities in tabular form such as local gene expressions retrieved from the Allen human brain atlas [33, 1]. For anatomical assignment, *siibra* computes correlation and overlap measures between the input image and brain area annotations in a reference atlas. In many cases it is advisable to split such feature maps into cluster components before computing the assignment (Figure 3 C). The given example uses hierarchical density-based clustering [13] of points sampled from the input image, based on an implementation in scikit-learn (RRID:SCR_002577) which is available through the `PointSet` class in *siibra-python*.^1^

As a result, for a given region of interest and a selected parcellation terminology, *siibra* implements two assessments. The first is an anatomical assignment of the region of interest, represented by a list of associated brain areas qualified by means of probabilities, spatial overlap or correlation, and their possible probabilistic maps in the corresponding coordinate space. The second is a multimodal profile, represented by a compilation of data features which characterize the region of interest. To support the FAIR principles of data organization, the underlying data elements are exposed in interoperable community formats and explicitly linked to the resource from which the information is retrieved. For example, tabular data are exposed as pandas DataFrames for optimal compatibility with common statistics and spreadsheet tools, while 3D images are exposed as NIfTI image objects which are supported by almost any neuroscience application.

In addition to the shown programmatic solution, *siibra-explorer* offers an interactive way to carry out a similar workflow using only a web browser, documented at https://siibra-explorer.readthedocs.io/en/stable/advanced/superimposing_local_files/. This is possible by drag-and-drop of an image volume from the user’s local computer onto the browser window. Assuming the volume is provided in NIfTI format and pre-aligned to the selected reference coordinate system, *siibra-explorer* will display it as an adjustable semitransparent overlay on top of the selected reference atlas, allowing direct visual assessment of overlap between brain areas and peaks or clusters depicted in the input image. Relevant data features can then be selected and downloaded from the side panel of *siibra-explorer*. The principle is visualized in a screencast video included in the documentation of *siibra-explorer*. However, the reproducibility of this viewer-based solution depends on the local environment, and is not supported per se in *siibra*.

### Multimodal comparison of brain areas

A common use case for the aforementioned anatomical assignment and multimodal profiling is to compare architectural features in different regions of interest. We demonstrate the principle by characterizing two cortical areas from the Julich-Brain atlas in terms of cell and neurotransmitter densities, gene expressions, and functional connectivity. As an example, data features of area 44 of Broca’s region (IFG) were compared with those of the primary visual cortex (hOc1/BA17/V1), both in the left hemisphere. The same principle can be applied to other brain areas and reference atlases, as well as to user-defined regions of interest. Retrieval of all information is implemented using *siibra-python* in a Python notebook included in the documentation. The feature plots displayed in Figure 4 allow for several interesting observations:

1. The two regions show distinct cortical architecture, revealed by different cell body distributions in cortical patches extracted from 1*µm* resolution scans of the BigBrain (Figure 4 A). Anatomically guided extraction of image data is covered in more detail below.
2. Average cell body densities across cortical layers differ between the areas, with overall higher values in the visual cortex, in particular the granular layers II and IV (Figure 4 B, top).
3. Receptor densities for selected monogenetic receptors (M1, M2, M3, D1, 5-HT1A, 5-HT2) show many similarities between area 44 and area hOc1, but also some notable differences such as a notably higher density of the M2 receptor in hOc1 (Figure 4 B, middle in hOc1). These receptors represent a subset of up to 19 available receptors, here reduced to those encoded by a single gene.
4. The expression patterns of genes coding for the selected monogenetic receptors (Figure 4 B, bottom) match well to the respective receptor densities (Figure 4 B, middle), which is in line with earlier studies highlighting such covariation between the modalities [71]. Gene expressions were retrieved by *siibra* from the Allen human brain atlas API [33, 1].
5. Functional connectivity profiles extracted from the HCP data [67] show distinct characteristic relationships for the two regions. For V1, we see strong connectivity into the hierarchy of higher order visual areas, following the organisation of the dorsal and ventral streams. For area 44 left, we observe strong temporal correlation with 44 right and 45, which together build the Broca region, as well as to the frontal pole, in correspondence with the known anatomy of the *arcuate fasciculus* [15].

**Figure 4.**
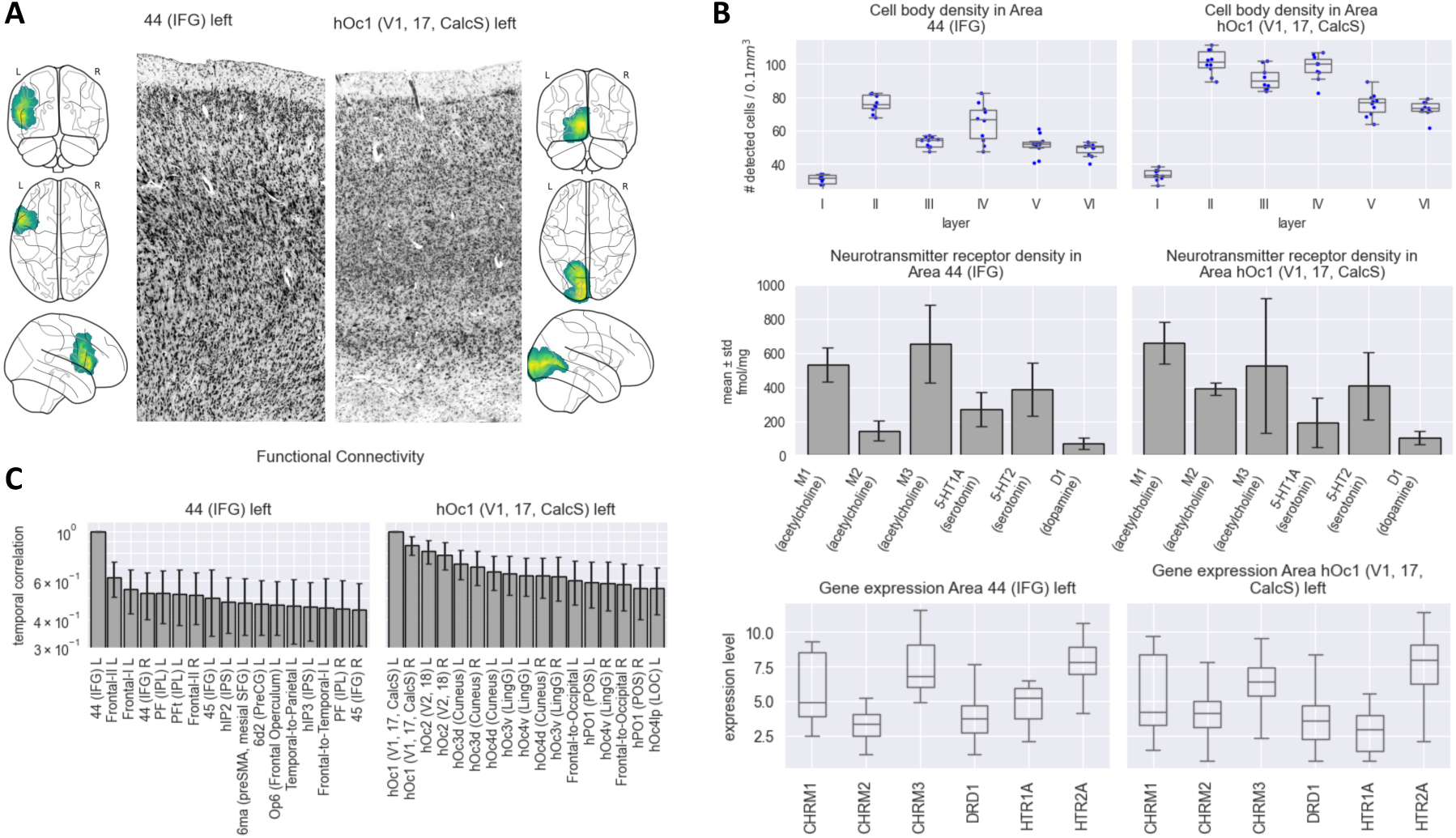
Multimodal comparison of two cortical brain areas. The figure shows maps and multimodal regional measurements obtained using *siibra-python* for language area IFG 44 and primary visual region hOc1, as defined in the cytoarchitectonic reference atlas [5]. **A**: Probabilistic maps in MNI asymmetric space, together with representative cortical image patches at 1*µm* resolution that were automatically extracted from scans of BigBrain sections [62, image contrast slightly increased] (cf. Figure 5). The image patches display clearly the differences in laminar structure of the two regions. **B**: Average densities of a selection of monogenetic neurotransmitter receptors [58] and cell bodies [22, 21], as well as expressions of a selection of genes coding for these receptors [33, 1]. **C**: Functional connectivity profiles referring to temporal correlation of fMRI time series of several hundred subjects from the Human Connectome Project [25, 67]. We show the strongest connections per brain area for the average connectivity patterns. The Figure can be reproduced using the notebook available at https://siibra-python.readthedocs.io/en/v1/examples/tutorials/2025-paper-fig4.html.

Here, we focus on summarizing observations to demonstrate key software functionalities, and refrain from delving into a deeper neuroscientific discussion, which would require additional analyses and statistical tests.

### Anatomically guided reproducible extraction of microscopy data

Besides retrieving of precomputed data features, *siibra* allows dynamic extraction of high-resolution image data from cloud image resources. For example, image intensities in the BigBrain model [4] can be interpreted as optical cell densities and thus provide a resource for studying basic aspects of cytoarchitecture. At an isotropic voxel resolution of 20 *×* 20 *×* 20*µm*, the BigBrain constitutes almost 1 TByte of image data, making it infeasible for many users to download and process the complete dataset. Using *siibra-explorer*, BigBrain can be viewed at full resolution by zooming in with the mouse wheel, such as the example region of interest bookmarked at https://atlases.ebrains.eu/viewer/go/bigbrain_precs_example. By hiding the parcellation map,^2^ the image data can be observed at full resolution (cf. Figure 2C).

However, interactive viewing of image data alone is not the preferred approach for performing reproducible studies and does not scale to large experiments that leverage the potential of big data resources. For this purpose, *siibra-python* can identify and fetch image data from cloud resources in an efficient and reproducible way. The underlying strategy is to specify a volume of interest using a bounding box or selecting a brain area from a reference atlas, then use *siibra-python* to identify relevant image datasets and extract the image parts intersecting the volume of interest. This process is facilitated by automatic handling of different coordinate spaces for the query, allowing fetching of data from multiple coordinate spaces using customized resolutions and fields of view, with automatic computation of the spatial metadata arising from this extraction process. The example in Figure 5 demonstrates how a cortical patch at 1*µm* resolution is automatically identified and fetched by selecting a cytoarchitectonic brain area. The resulting data is provided as a NIfTI image annotated with the appropriate transformation matrix into the BigBrain reference coordinate system. As such the fetched data is suitable for downstream analysis tasks such as cell segmentation, and for combination with other image data referring to supported reference coordinate systems.

**Figure 5.**
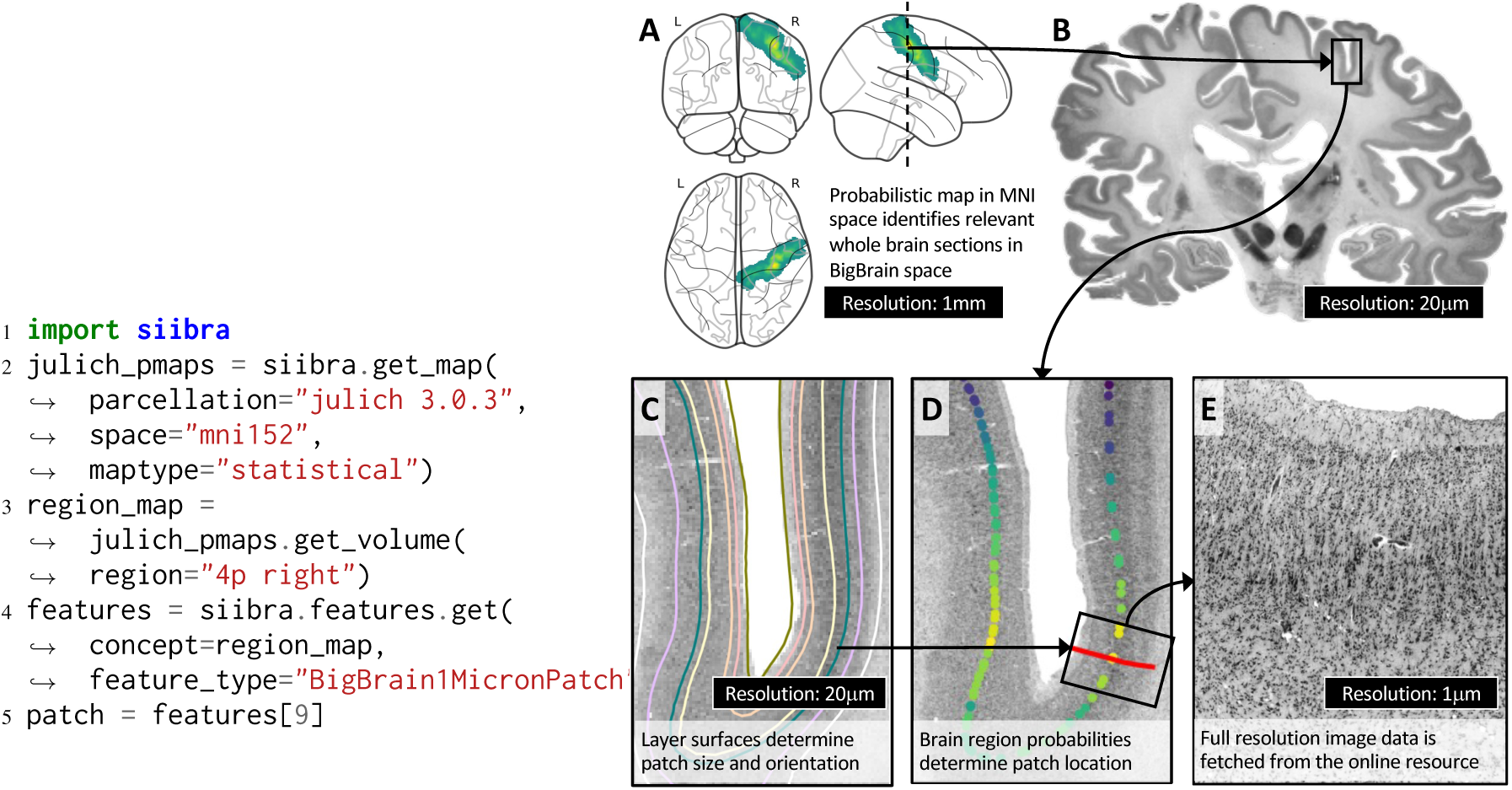
Anatomically guided reproducible extraction of full resolution image data from cloud resources. *siibra-python* allows implementing reproducible workflows for sampling microscopy data from anatomically defined regions of interest. The code example on the left retrieves the probabilistic map of area 4p in the right motor cortex from the cytoarchitectonic atlas [5] (A; ll. 2-3), and uses it to sample relevant cortical patches from high-resolution scans of whole-brain tissue sections in BigBrain space [4, 62] (B; l. 4). To define an oriented cortical image patch, *siibra* automatically loads and intersects cortical layer surface meshes [68] with each relevant section image (C). It then finds points on the intersected layer IV surface with high relevance according to the probability map of the motor area (D). The different coordinate systems in this process are automatically handled using nonlinear transformations as detailed in the “Methods” section. For a given point on the layer IV surface, *siibra* finds the closest 3D cortical profile in the layer maps to obtain information about orientation and thickness of the cortex at the chosen position. The 3D profile is projected to the respective image plane and used to define a rectangular patch. The full resolution image data inside the cortical patch is then fetched from the underlying cloud resource (E; l. 5). The resulting patch can be readily used to extract cell body instances and compute quantitative density measures using state-of-the-art instance segmentation models, resulting in cell densities as published in a recent data collection (Figure 4B, e.g. [22]). *The Figure can be reproduced using the notebook available at* https://siibra-python.readthedocs.io/en/v1/examples/tutorials/2025-paper-fig5.html.

### Expansion with new contents

By maintaining a strict separation from the software, the *siibra* tool suite allows the content of the Multi- level Human Brain Atlas to continue to expand. In particular, any terminology, parcellation map, reference template or data feature exposed by *siibra* points to an external resource which contains the actual data (Figure 1A). This can be a file on a public data repository such as EBRAINS, a segment of a cloud image volume, a local file provided by the user, or data that is dynamically retrieved from a cloud service. We distinguish *foundational content*, defined as json specifications in a configuration, from dynamically retrieved content obtained through *siibra*’s *live queries*, and *local content* provided by users on their own system. These different routes are reflected in the software design (Figure 6 A-C) and result in the following alternative ways for expanding the Multilevel Human Brain Atlas:

1. New foundational contents are added to *siibra*’s configuration by creating specification files. This allows flexible addition of different object types from any accessible data resource. For adding specifications to the default configuration of the Multilevel Human Brain Atlas, a change request can be submitted at https://github.com/FZJ-INM1-BDA/siibra-configurations. Local installations of *siibra-python* can use custom configuration repositories, which allow to clone an existing configuration to a local computer add new specification files directly.^3^
2. Dynamic content can be made discoverable in *siibra* live queries by sharing it on a supported cloud service. For example, when releasing a public dataset on EBRAINS, information about its anatomical location will be modelled according to the openMINDS SANDS metadata model (RRID:SCR_023498) and stored in the EBRAINS Knowledge Graph (RRID:SCR_017612). This allows *siibra*’s EBRAINS live query to match it with atlas elements and include it automatically. To connect new cloud services, *siibra-python* can be extended with new live queries.
3. Local content, including both reference atlases and data features, can be added adhoc in *siibra-python* using various utility functions, such as in Figure 5C. For example, users may add a custom parcellation map or an image volume to query data features, demonstrated for the AICHA atlas [38] in the code example available at https://siibra-python.readthedocs.io/en/v1/examples/02_maps_and_templates/007_adding_custom_parcellation.html. In *siibra-explorer*, user supplied NIfTI volumes can be visualized on top of existing reference atlases by drag-and-drop onto the browser window, as illustrated in the documentation at https://siibra-explorer.readthedocs.io/en/stable/advanced/superimposing_local_files/. Assuming it has been pre-aligned to the respective reference coordinate space, the file is co-visualized in the browser while solely remaining on the user’s local system.

**Figure 6.**
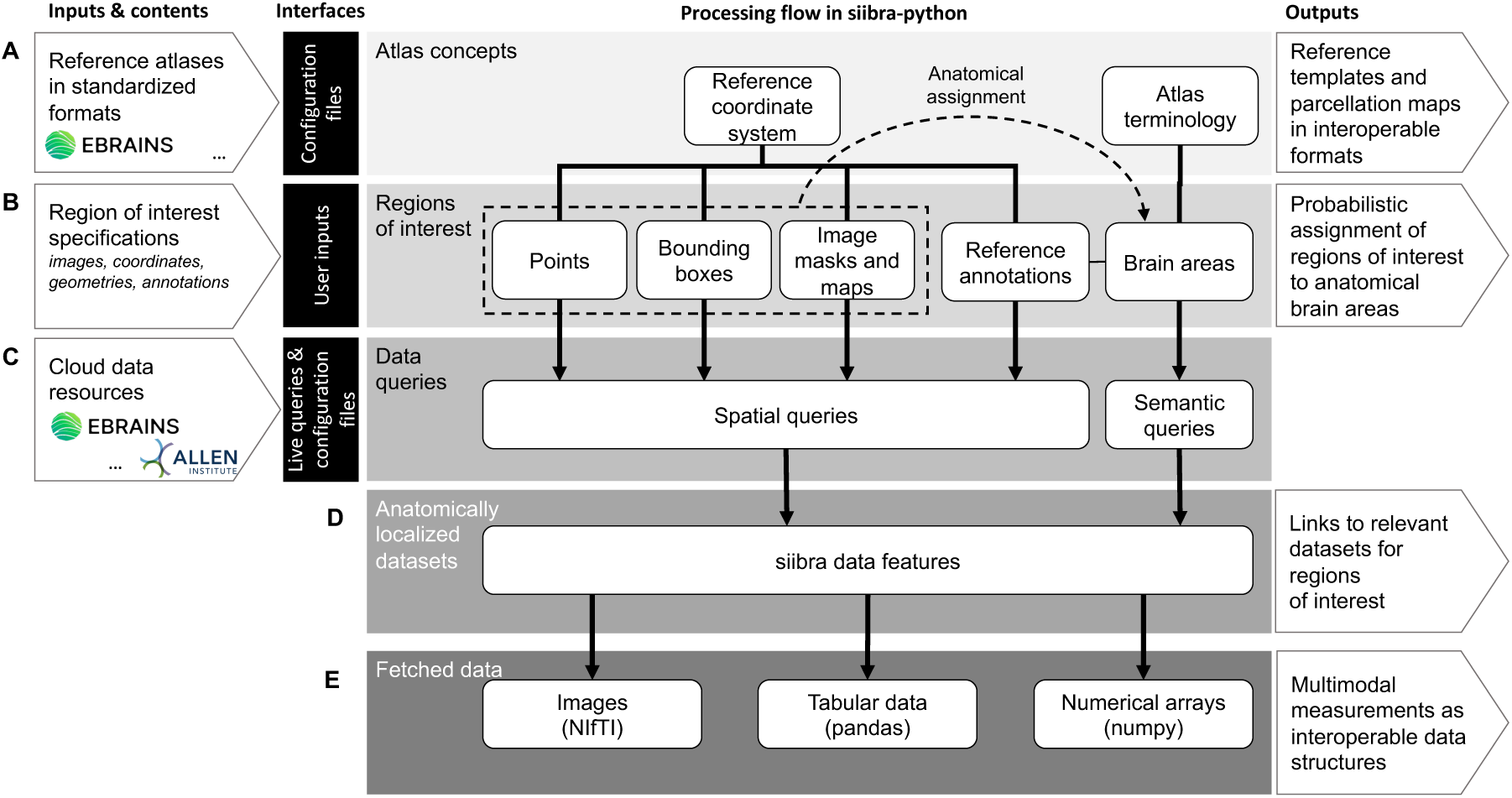
High-level illustration of *siibra-python*’s system architecture. The *siibra-python* library models elements of reference atlases and spatial locations to enable assignment of regions of interest to brain areas, and implements queries for retrieving anatomically localized data features from cloud resources. **A**: Atlas elements include reference coordinate spaces and parcellations with corresponding terminologies, which are stored in standardized form in data repositories such as EBRAINS and linked to *siibra* through configuration files. The library provides efficient and reproducible retrieval of maps and templates at different spatial resolutions in interoperable formats such as NIfTI. **B**: Users can specify regions of interest by providing images, coordinates, geometric shapes or textual annotations, which *siibra* assigns to brain areas via probabilistic reference maps. Resulting assignments can be accessed and reused as pandas DataFrames. **C**: Regions of interest and brain areas can then be used to run queries to cloud repositories to obtain customized multimodal profilings. Foundational datasets are linked to *siibra* in the form of configuration files, and complemented with dynamic content resulting from API calls (“live queries”), currently including interfaces to EBRAINS and the Allen microarray data for gene expressions in the human brain [33, 1]. **D**: Data queries result in *data features*, which can be considered as enriched links between anatomical locations in *siibra* and files published on the respective online resource. The links can be accessed in the form of URLs and DOIs together with a qualification of their anatomical anchoring to the atlas. **E**: By fetching from a data feature, *siibra* provides access to the underlying data in interoperable formats such as NIfTI images, pandas DataFrames, or numpy arrays for simple reuse. Following a lazy loading scheme, data is retrieved from the cloud resource only when required, and then cached for future requests.

In addition to the Multilevel Human Brain Atlas presented here, *siibra* can be used for brains of other species without advanced programming skills. It has been successfully applied to set up the macaque brain atlas [7, 56, 61] available at https://atlases.ebrains.eu/viewer/go/mebrains, the Waxholm Space Atlas of the Sprague Dawley Rat Brain [41, 57, 59, 40] available at https://atlases.ebrains.eu/viewer/go/rat, and the Allen mouse brain atlas [69] available at https://atlases.ebrains.eu/viewer/go/mouse. A minimal configuration for a new atlas requires three json specifications, namely 1) a terminology of brain areas, 2) a reference coordinate system with an anatomical template in volume or surface form, and 3) a parcellation map in volume or surface form defined in the same coordinate system. This implies to host at least two image volumes or surface meshes on a web-accessible resource.

## Discussion

The development of a Multilevel Human Brain Atlas atlas covering the scales from cells and fibres up to the macrostructure of the entire organ requires novel conceptual and technical solutions. To exploit the full potential of large-scale, high-resolution measurements from modern imaging techniques, the operationalization of such a system inevitably requires instruments for handling big data resources.

We have solved this challenge by using the *siibra* tool suite to implement an openly accessible and inter-operable Multilevel Human Brain Atlas. The tool suite facilitated a systematic technical realization of the atlas by linking complex multimodal datasets with reference parcellations in different coordinate systems, including full-resolution image data from microscopic experiments. We use cytoarchitecture as a fundamental principle of brain organisation in the atlas, because it provides an adequate spatial scale and represents a common reference modality for neuroscientific findings. The use of probabilistic maps for assigning regions of interest to brain areas defined in reference atlases is encouraged, as a statistical treatment is crucial for modeling common uncertainties of in vivo analyses. Being designed as a distributed system that can connect to a multitude of configurable cloud resources, *siibra* embraces the rapid growth in high-throughput imaging techniques and increasing availability of big data repositories. The tool suite is well integrated and supported by the EBRAINS research infrastructure which allowed us to connect it with a broad compilation of datasets, tools and online services. At the time of writing, the Multilevel Human Brain Atlas provides access to more than seven terabytes of multimodal datasets in the form of 2D and 3D images, tables and numerical arrays, but its design allows for continuous expansion with new instances of reference atlases and data features.

The *siibra* tool suite offers accessibility to user groups with different programming skills: Non-programmers can engage interactively via a web browser, computationally adept neuroscientists can utilize scripting and workflow construction in Python, and professional programmers can leverage a web API for application development. The versatility therefore spans from neuroimaging and artificial intelligence to computational neuroscience and clinical applications. This has been demonstrated in first studies applying *siibra* successfully, e.g. for modeling whole brain dynamics with regional heterogeneity [64] and investigating structure-function relationships [72].

In either form, *siibra* assumes minimal technical requirements. The online viewer can be accessed with common web browsers, including mobile devices, and co-display user-supplied NIfTI files with atlas content without uploading them. The Python client can be installed with established package managers from https://pypi.org/project/siibra and provides a broad range of documented code examples at https://siibra-python.readthedocs.io/en/v1/examples.html. Because content is separated from software, installation of the Python client requires only minimal storage space. Following a lazy data fetching strategy, *siibra* downloads only requested data elements to gradually build up a configurable local data cache according to user needs. For operating the system in commercial or clinical environments with restricted network policies, *siibra* can be setup for offline use by cloning the configuration to a local computer, pre-populating the cache explicitly and deploying the API and viewer components as local network services.

The development of the Multilevel Human Brain Atlas using *siibra* was closely integrated with ongoing community efforts in metadata standardization, resulting in foundational atlas content being curated according to the openMINDS metadata framework (RRID:SCR_023173), and fundamental design principles in *siibra* following the AtOM ontology model [42]. The current version of *siibra* puts a focus on foundational data resources established in EBRAINS by the Human Brain Project [3], but it is designed from ground up to interface more broadly with a growing portfolio of neuroscience community resources. Upcoming releases of *siibra* will link with recently proposed repositories for collecting file-based MRI-scale reference templates and parcellations [60, 16, 44, 17] and online resources for additional data modalities such as human cell morphologies [65] and functional measurements [52]. The integration of such data resources on the basis of *siibra* live queries is much facilitated by the growing adoption of the BIDS standard [31] and its recently proposed extensions for other data modalities (e.g. [11]).

As big data and AI increasingly become central pillars of neuroscientific research [23, 35], we face the challenge of how to involve researchers in complex reasoning and discovery processes. The combination of big data visualization (in *siibra-explorer*) and systematic data processing (in *siibra-python*) provides new opportunities for designing visual analytics (VA) workflows [66, 18, 48]. These aim to combine purely data-driven exploration with human cognitive, perceptual and reasoning abilities by building visually supported loops of data extraction, hypothesis building, and inference. With the advent of AI and big data, visual analytics is expected to gain increasing interest in the neuroscience community in the future. Software solutions such as *siibra* are key enablers of such developments.

From a technical perspective, *siibra* covers core principles of data aggregation systems [12, 14] by addressing the collection, processing, and combination of data from multiple sources to better enable downstream tasks such as hypothesis generation, analytics, modeling and inference, and by solving problems such as redundancy, inconsistencies, and varying data formats. In brain research, addressing these informatics challenges is crucial for harnessing data in large-scale, data-driven models like AI foundation models, anatomically accurate simulations, and digital twins, paving the way for new research questions to be explored. Of course, data aggregation systems play a crucial field in other fields. Although *siibra* was implemented for neuroscientists in the first place, the key aspects of its implementation are not specific to this field. With moderate adaptions of the code, it could serve as a basis for building atlases of other organs such as the heart or kidney, and more generally for data aggregation of systems with a strong 3D topographical organisation which are described by images and tabular content at multiple scales.

In conclusion, the *siibra* tool suite introduces a highly configurable informatics framework for building multilevel atlases of different species and organs, which 1) provides three levels of operability with an API, a web viewer and a python library; 2) enables the handling of large image datasets (“big data” capabilities); 3) provides interoperability with distributed data resources and community tools - thus making atlas data actionable; 4) supports the FAIR principles; and 5) is designed in a flexible way to address the continuous expansion with content. It is supported by the EBRAINS research infrastructure, and open for contributions from the community.

## Methods

In the following we describe relevant methods and technical principles applied in the *siibra* tool suite, on which the presented results are based. These include modularization of the software, separation of code from contents, and strategies that enable the use of large image datasets and locations in different reference coordinate systems.

### Modeling spatial entities in different reference coordinate systems

In line with community ideas [2, 42], *siibra* distinguishes semantic atlas elements such as reference coordinate systems and terminologies from their spatial representations as annotations sets or reference templates. This is a prerequisite for using the same anatomical concepts in processes operating at different spatial scales or topological layouts. For example, the same atlas terminology can be realized by annotation sets in different reference coordinate systems, thereby resulting in multiple representations of the same brain area with different coordinate system, spatial resolution or data format. Similarly, a reference coordinate system can be represented by different reference datasets (i.e., templates) with specific resolution and representation of the underlying anatomy as an image volume or surface.

In general, points in different reference coordinate systems are related by nonlinear coordinate transformations. This association is not unique and generally inaccurate, since different mathematical models and optimization strategies for the transformation result in different solutions. For the Multilevel Human Brain Atlas, we used diffeomorphisms constrained with cortical folding patterns [50, 45] to obtain a precise alignment between reference coordinate systems in regions where sulci can be used as landmarks. These transformations are exposed through a web service (https://github.com/HumanBrainProject/hbp-spatial-backend) which *siibra* uses to transform coordinates between reference coordinate systems when querying data features or comparing spatial objects. All spatial entities in *siibra-python* reflect instances of *locations* such as points, point clouds or bounding boxes (Figure 6). Examples of these three types of instances are centroids of brain areas, bounding boxes of volumes of interest, contact points of electrodes, peaks of fMRI activations maps, or the currently selected view in *siibra-explorer*. Since any such location object is uniquely associated with a reference coordinate system, coordinate projections can be carried out automatically, thus allowing to apply most spatial operations in *siibra* safely to objects from different reference coordinate spaces. This principle has been demonstrated in the results for cortical image patch sampling, where the bounding box of a brain area in MNI space is automatically transferred to BigBrain space for finding high-resolution image data (Figure 5) and a set of vertices in BigBrain space is evaluated by a probability map in MNI space.

### Linking data features to atlas elements

The localization of data features with respect to atlas elements can take various forms with different levels of precision. In a human brain atlas, this is particularly influenced by the high inter-subject variability resulting in significant uncertainties when relating spatial locations measured in different brains. The most accurate localization is possible for measurements obtained in the same brain, where e.g. image registration can provide adequate alignments. For different brains, an accurate correspondence can be achieved if the data allows reliable histological identification of the underlying brain area. This principle is used extensively for the Multilevel Human Brain Atlas: Cytoarchitectonic maps are being used as a preferred reference to localize foundational data features such as neurotransmitter receptor and cell densities according to cytoarchitectonic criteria of the tissue samples. This provides an *intrinsic* definition of anatomical locations, directly grounded in properties contained in the measurement, and therefore independently verifiable. For other datasets such as in vivo neuroimaging or physiological recordings, the assignment often has to rely on projections of coordinates. This can be seen as an *extrinsic* transfer of the localization to the data, which is not uniquely defined and less verifiable.

To capture different forms of anatomical correspondence between atlas elements and data features, *siibra* tags anatomical locations with semantic relationships to brain areas from a reference parcellation, with spatial locations such as points, lines, or bounding boxes in a reference coordinate space, or combinations thereof. As such it implements basic principles of the *Locare* workflow proposed for representing locations in 3D brain atlases [9]. The semantic or spatial location specifications can be further refined by certainty quantifications such as information about containedness and overlap of brain structures, accuracy of coordinates, or hierarchical dependencies between child and parent areas in an atlas terminology.

As a result, *siibra* can query many forms of data features in a unified fashion by providing an anatomical concept (such as a brain area or volume of interest) and a modality for the requested data features. When evaluating such a query, *siibra* uses all available information to compute a qualification for each matched data feature. This enables an adequate assessment of the relevance and accuracy of retrieved measurements with regard to a given analysis task. A practical example of this principle can be reproduced using the notebook available at https://siibra-python.readthedocs.io/en/v1/examples/03_data_features/000_matchings.html.

### Modular software architecture

The *siibra* tool suite consists of an interactive 3D web viewer (*siibra-explorer*), an HTTP API (*siibra-api*) and a Python library (*siibra-python*).^4^ Following a modular software design, these components share most of their functionality while addressing different user scenarios. The core functionality is implemented in *siibra-python*, a Python package suitable for scripting, interactive notebooks and development of computational workflows. The key elements of the system architecture are illustrated in Figure 6. It models various elements of reference atlases, coordinate-based locations, data features and queries; and provides functionalities for loading, matching and processing these entities. Since contents are managed separately from the software, efficient retrieval of data from cloud resources becomes an essential aspect of the package. Various data fetching mechanisms are implemented for files, multi-resolution images and metadata, following a lazy loading strategy to ensure that only actually requested elements are transferred. The package documentation is hosted at https://siibra-python.readthedocs.io and automatically updated with new releases. It contains a comprehensive collection of documented and tested code examples which demonstrate typical use cases, including the executable codes to reproduce the figures in our results. These are complemented by a growing collection of more comprehensive tutorial notebooks available at https://github.com/FZJ-INM1-BDA/siibra-tutorials, developed for workshops and user trainings.

The functionalities of *siibra-python* are exposed as a RESTful API for application development (*siibra-api*). This HTTP interface focuses on the processing of HTTP requests and efficient serialization of *siibra* objects and method calls using FastAPI as a web framework, and implements type-hinting via OpenAPI. A production instance is deployed, documented and openly accessible for application developers at https://siibra-api.apps.ebrains.eu/v3_0/docs.

Finally, the web viewer *siibra-explorer* implements an interactive user interface on top of *siibra-api*— focusing on visualization and user interactions while reusing the underlying functionalities from *siibra-python*. It uses Angular as a reactive programming framework, neuroglancer [49] as a visualization backbone for very large multi-resolution volumetric images, and ThreeJS for rendering of brain surfaces. The surface view is automatically activated when selecting the freesurfer space, and provides white matter, pial and inflated surface representations. The viewer can be accessed from common web browsers, including mobile devices with touch inputs such iOS or Android tablets.

The production instance of the viewer provides access to the Multilevel Human Brain Atlas presented in this paper and is available at https://atlases.ebrains.eu/viewer. To track the community interest in the atlas, we record the numbers of unique returning visitors per month on this instance, which has been increasing over a timescale of 4 years from about 400 (summer 2020) to about 1300 (summer 2024).

Documentation for *siibra-explorer* is hosted at https://siibra-explorer.readthedocs.io/en/latest/. It provides a guided quick tour to help users getting started, which is automatically launched on first use and can later be accessed via the help menu.

To keep the development of *siibra-explorer* focused on its core requirements while still allowing flexible addition of functionalities, the viewer supports the implementation of plugins similar to the add-on mechanisms offered by modern web browsers. This has been used to create an interactive implementation of the JuGEx workflow [10], which can be accessed from the plugin menu when selecting the MNI 152 reference space. The plugin architecture has also been used successfully by researchers to integrate a workflow for learning gene regulatory networks with Bayesian networks [8].

### Separation of content from code

A fundamental design principle of *siibra* is separation of content from software—an important prerequisite for extensibility.

#### Specification files

The foundational content elements such as parcellation maps, reference templates and data features are connected to *siibra* through *specification files* maintained outside the codebase. These specifications follow a set of simple schemata maintained in the `config_schema` folder of *siibra-python*. Typically, a specification file defines 1) minimal metadata entries such as identifier, name, and modality, 2) pointers to an external metadata management system such as the EBRAINS Knowledge Graph providing extended metadata, 3) an anatomical reference such as a coordinate-based location or a brain area name from a reference atlas, and 4) URLs and format specifiers that identify the underlying data, such as image or tabular data files stored on a cloud resource. The specification of a basic image data feature, taken from the configuration at https://github.com/FZJ-INM1-BDA/siibra-configurations/tree/v1/features/images/sections/cellbody, is displayed in the Supplementary Information, Figure S1. While some specifications such as for parcellation maps are more comprehensive, it shows the typical style and organisation of the format. A more complete documentation of *siibra* specifications is available as part of the code at https://github.com/FZJ-INM1-BDA/siibra-python/tree/v1/config_schema.

#### Configurations

Foundational content specifications are collected into *configurations* by packaging json files into a zip container, git repository, or folder on a file system. Configurations can be hosted both on a local machine or on a cloud service. The default configuration of *siibra* provides the foundational content of the Multilevel Human Brain Atlas. It is maintained at https://github.com/FZJ-INM1-BDA/siibra-configurations, continuously updated by the maintainers and open for extension requests by the community. Local installations of *siibra* can use alternative or extended configuration repositories with customized content, ensuring well-defined maintenance, extension and customization of atlas contents.

#### Live Queries

Foundational content specified in configurations is extended by dynamic content from *siibra live queries*—software interfaces implemented in *siibra-python* for connecting to APIs of suitable cloud services and translating their responses into *siibra* specifications at runtime. Implementing live queries assumes external repositories to provide a stable API and to return sufficient metadata and common access URLs to generate specifications as described above. So far, *siibra* includes live queries to the EBRAINS Knowledge Graph (RRID:SCR_017612) for discovering anatomically anchored datasets, to the Allen Human Brain Atlas [33, 1] for accessing microarray data localized in MNI space, and to cloud image resources, e.g. [62], for sampling high-resolution image data as demonstrated in Figure 5.

### Support of the FAIR principles

Most foundational content in *siibra*’s default configuration points to curated datasets in EBRAINS, supporting the FAIR principles for data sharing [70]. Registration in the EBRAINS Knowledge Graph (RRID:SCR_017612) ensures *findability*, as it provides each content element an individual DOI, links it with rich metadata, and offers comprehensive faceted search interfaces. The search paths offered by the Knowledge Graph are further complemented by *siibra* in terms of more advanced anatomical navigation and atlas-guided processing. *Accessibility* of rich metadata is provided for atlas content through their DOIs, which are exposed by *siibra* and allow direct online access to original dataset pages including comprehensive data descriptors. The *siibra* tool suite adds additional entry points for retrieval by forwarding the metadata, offering download links in *siibra-explorer*, and implementing efficient data fetching capabilities in *siibra-python*. By promoting the openMINDS metadata framework (RRID:SCR_023173) used by the EBRAINS Knowledge Graph (RRID:SCR_017612), *siibra* further supports *interoperability and reusability*. Instead of establishing a different knowledge representation, *siibra* specification schemas only serve as a summary and reference pointer to the established representations and terminologies in EBRAINS. By explicitly subjecting the content of each foundational atlas element to the professional data curation offered by EBRAINS, a comprehensive description of the content with attributes accepted by the community is ensured. Interoperability is further optimized by exposing the retrieved contents from the underlying dataset in highly interoperable formats. For example, in *siibra-python*, fetched image data is exposed as Nifti1Image image objects via NiBabel (RRID:SCR_002498), fetched tabular data as Pandas (RRID:SCR_018214) DataFrames, and more general numerical data such as vertex lists as NumPy (RRID:SCR_008633) arrays (Figure 3). This makes it straightforward to use the data with popular tools such as e.g. NiLearn (RRID:SCR_001362), itksnap, Napari (RRID:SCR_022765) and MatPlotLib (RRID:SCR_008624), and integrate it in common analysis workflows.

### Unified access to small and big image volumes

A central design goal of *siibra* is to unify access to small and “big” image data, e.g. to handle images in the range of a few hundred Megabytes up to Terabyte scale in a consistent fashion while hiding the complexity of their different access paths as cloud volumes or downloadable files. This has been realized by supporting both conventional local and web-hosted NIfTI files as well as multi-resolution chunk formats such as the neuroglancer precomputed tile format through the same user-level functionality. The heterogeneity of these data types is hidden behind a common interface in *siibra-python* which provides image data in NIfTI format with appropriate spatial metadata independently of its original format and origin. When accessing content elements with conventional “small” 3D image data, *siibra* will load and forward the original image data in unmodified form. When fetching from a large multi-resolution image resource, *siibra* will either return an appropriately downscaled version or require to specify a region of interest that results in a feasible download size (Figure 5E). In any case, the involved cropping or scaling will be reflected in the spatial metadata of the resulting NIfTI object, so that the fetched data remains appropriately localized in its original reference space. This way, *siibra* can convert large whole-brain image resources stored on cloud resources into actionable data structures as required by the user. This principle can be reproduced using the notebook available at https://siibra-python.readthedocs.io/en/v1/examples/02_maps_and_templates/004_access_bigbrain.html.

### Memory-efficient representation of sparse regional maps

Statistical maps associate 3D points in a reference coordinate space with continuous weights reflecting the likelihood of a brain area being present at that location [2]. Processing such statistical parcellation maps e.g. for probabilistic assignment (Figure 3) thus might imply retrieving and loading one image volume per brain area. For Julich-Brain this amounts to loading approximately 300 3D volumes in MNI space, which results in long download times for the initial data fetching and may quickly exceed the working memory capacity of standard computers. To avoid these problems, *siibra* implements an alternative data structure to minimize memory consumption. The structure exploits that statistical region maps often encode only a small fraction of the brain volume, thus mostly containing zeros and effectively forming an extremely sparse 4D array over all brain areas. The main idea is to replace these sparsely populated 3D volumes by an index which stores only the nonzero weights. This is modelled by two data structures, which may be understood as a “spatial” dictionary with coordinate-based keys (see Supplementary Information; Figure S2): The first structure is a single 3D array matching the original voxel dimensions of the maps. This array structure allows efficient spatial queries with bounding boxes or image masks. Elements of the array are zero if the corresponding voxel is not associated with any brain area in the parcellation, or hold a numerical key otherwise. The numerical keys are indices to the second structure, a list containing the statistical weights of the corresponding voxel statistical maps concerned. The structure reimplements all functions of conventional parcellation maps, including data fetching and assignment of coordinates to brain areas.

## Acknowledgments

We thank P. Chervakov for developing a wrapper (“nehuba”) around the neuroglancer viewer for better integration as a 3D brain viewer; V. Marcenko for contributing to the development of early prototypes of *siibra-python* and *siibra-api*; D. Gogshelidze for implementing the first version of a widget for connectivity features in *siibra-explorer*; L. Köhnen for contributing to the development of an early prototype of *siibra-python*; L. Zehl for guidance on connecting openMINDS to *siibra*; J. Thoennissen for testing *siibra-python* and *siibra-explorer*; J. Fousek for valuable feedback on the *siibra-python* interface; M. Hanke for insightful discussions about modeling data elements in the software architecture; S. Wenzel and K. Lothmann for organizing numerous workshops, tutorials and trainings with *siibra* users; A. C. Evans, R. N. Kooijmans, C. Lepage, K. Lothmann, M. Puchades, S. Ruland, and P.-J. Toussaint for constructive feedback regarding the user interface of *siibra-explorer*; C. Rorden for helpful discussions regarding the technical representations of maps; D. Rivière, J. Chavas and J. Lebenberg for providing the sulcus-based diffeomorphic transformations between reference spaces and M. Axer, N. Palomero-Gallagher, K. Wagstyl and C. Poupon for sharing their deep knowledge about foundational data resources such as 3D-PLI images, neurotransmitter receptor densities, cortical layer segmentations, and ultra-highfield MRI volumes.

## Code availability

Source codes for *siibra* are available on GitHub (https://github.com/FZJ-INM1-BDA/siibra-python; https://github.com/FZJ-INM1-BDA/siibra-explorer; https://github.com/FZJ-INM1-BDA/siibra-api) and provided under the Apache 2.0 license (http://www.apache.org/licenses/). We have integrated all components of *siibra* with Zenodo, which generates unique digital object identifiers (DOIs) for each new release of the tool suite. An installation of the interactive viewer is publicly accessible at https://atlases.ebrains.eu/viewer as a core component of the brain atlas services in the research infrastructure EBRAINS. The web API can be accessed at https://siibra-api-stable.apps.hbp.eu/v3_0/redoc. Researchers can install *siibra-python* through the Python Package Index from https://pypi.org/project/siibra. Comprehensive online documentation, including executable codes to reproduce all Figures can be accessed via https://siibra-python.readthedocs.io/en/v1. More detailed tutorials notebooks are provided at https://github.com/FZJ-INM1-BDA/siibra-tutorials.

## Availability of data and materials

All data used in this work are publicly available via the URLs and references provided with the respective figures. With few exceptions, contents are published as curated datasets with permanent identifiers on the EBRAINS platform (https://ebrains.eu). The current foundational content is defined and maintained at https://github.com/FZJ-INM1-BDA/siibra-configurations/tree/v1. An overview is provided in the Supplementary Information, Table S3.

## Authors’ contributions

T.D. and K.A. conceived the idea of implementing a software tool suite to realize a multilevel atlas of the human brain. T.D. conceptualized the scope of the tool suite and led the overall development. X.G. is the main developer of *siibra-explorer* and *siibra-api*. T.D., X.G. and A.S. are the core developers of the *siibra-python* library. S.B. and C.S. provided substantial inputs to the conceptualization and numerous critical reviews regarding the usability of the tool suite and foundational content. J.-F.M. and Y.L. provided non-linear transformations between coordinate systems and the implementation of the corresponding backend service for coordinate warping. V.J. provided substantial guidance for making the tool suite and foundational contents usable for modeling and simulation, particularly integrating it into modeling workflows and selecting relevant foundational content. J.G.B. and T.B.L. provided continuous guidance regarding the compatibility of the tool suite with openMINDS and AtOM, its suitability as a core service component in EBRAINS, as well as its utility for building atlases of brains from other species. T.D. and K.A. wrote the manuscript with contributions from all authors.

---

**Figure S1.**
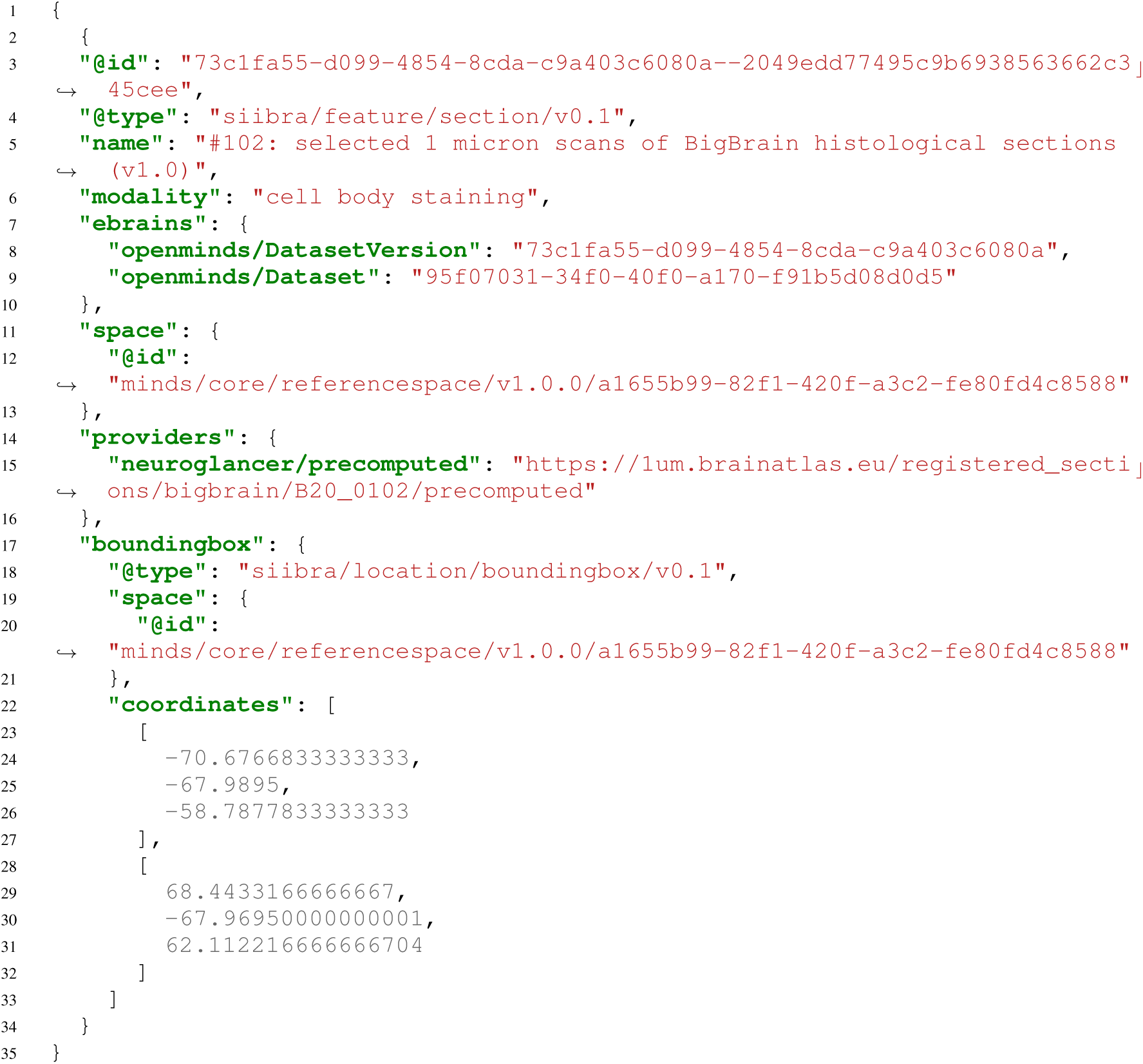
Specification of a typical image data feature. The specification is given as a json file and describes a 2D section anchored to the BigBrain reference space [62]. The original file is part of *siibra*’s default configuration available at https://github.com/FZJ-INM1-BDA/siibra-configurations/tree/v1/features/images/sections/cellbody. It follows the schema `section/v0.1` defining text fields for dataset name and modality, as well as URLs to relevant openMINDS metadata entities of the corresponding dataset in EBRAINS. The actual content is identified by the URL to the multi-resolution image data in neuroglancer precomputed format, and associated with the BigBrain reference space using a unique identifier. Using this information, *siibra* is able to extract detailed metadata and access the full resolution image data as demonstrated in Figure 2D-F and Figure 5E.

**Figure S2.**
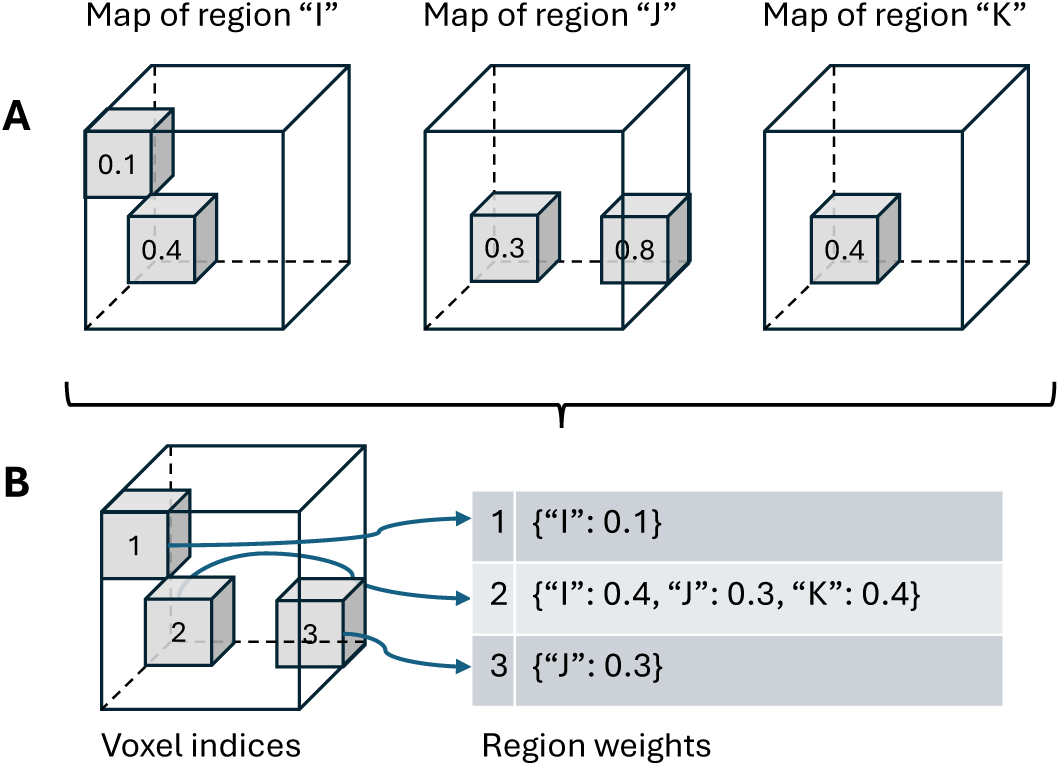
Representation of sparse statistical parcellation maps used in *siibra*. **A**: Three hypothetical statistical maps for different brain areas “I”, “J” and “K”, represented as simplistic arrays of 3 *×* 3 *×* 3 voxels with only a few nonzero weights each. **B**: The statistical maps are converted into one single array storing indices to all “occupied” voxel coordinates, together with a list of the map weights associated with these voxels. The weights for a given voxel are stored in the list as a dictionary which maps from the corresponding brain area to the weight.

**Table S3.** Snapshot of foundational content in the Multilevel Human Brain Atlas. The table summarizes the set of foundational data specified in the default *siibra* configuration of the Multilevel Human Brain Atlas at the time of writing, including maps, templates and multimodal data features. The amount of data features that can actually be accessed is larger due to dynamic content obtained through live queries (Figure 1A) as well as continuous extension of the foundational content in *siibra*’s default configuration. The reported version of the configuration is available at https://github.com/FZJ-INM1-BDA/siibra-configurations/tree/v1.

| Category | Modality/Type | Source | Description | ≈ GiB |
| --- | --- | --- | --- | --- |
| maps | cytoarchitecture | [5] | Julich-Brain probabilistic maps of cortical and subcortical areas | 3 |
|  | fibre architecture | [32] | Probabilistic maps of superficial and white matter bundles | 1 |
|  | function | [19] | statistical maps of functional modules | 1 |
|  | cortical layers | [68] | BigBrain maps of isocortical layers | 1 |
| templates | Microscopy | [4] | BigBrain histological 3D reconstruction | 260 |
|  | MRI | [34] | MNI Colin 27 | 1 |
|  | MRI | [29] | ICBM 2009c nonl. asym. | 1 |
|  | Surface | [27] | Freesurfer fsaverage and fsaverage6 | 1 |
| cellular features | Cortical cell distributions | e.g. [22] | Image patches at $1\mu m$ resolution with cell segmentations & layer annotations | 10 |
|  | Cortical staining profile | [68] | 163828 profiles in left BigBrain hemisphere | 1 |
| | Cell-body stained whole brain section | [62, 4] | 145 coronal whole-brain sections at $1\mu m$ resolution | 4 500 |
| | Cell-body stained volume of interest | [36] | 2 volumes of interest in occipital lobe scanned at $1\mu m$ | 410 |
|  | Phase-contrast tomography | [26] | Reconstructed density of a hippocampal tissue volume | 61 |
|  | LSFM | [54] | Human brain cell census for BA 44/45 | 58 |
| molecular features | Receptor density fingerprint | [58, 73] | Average densities of up to 16 receptors in 42 brain regions | 1 |
|  | Cortical receptor density profile | [58, 73] | 107 profiles of 16 receptors across 7 brain regions | 1 |
| fibre architecture | 3D-PLI | [6] | Fibre orientation maps of a hippocampus from 3D Polarized Light Imaging | 2070 |
| | DTI | | Whole brain post-mortem fractional anisotropy and RGB maps at $200\mu m$ | 35 |
| macro-anatomy | Blockface | [6] | Blockface reconstruction of a human hippocampus | 1 |
|  | MRI | [6] | T2 weighted MRI of a human hippocampus | 1 |
| | MRI | | Ultra-highfield anatomical MRI of a post-mortem brain at $200\mu m$ and $150\mu m$ | 107 |
|  | MRI | [46] | Ultra-highfield anatomical MRI of human brainstem | 6 |
| connectivity matrices | Streamline counts | [39] | Streamline counts of 821 subjects from 2 cohorts averaged for 3 parcellations | 1 |
|  | Streamline lengths | [39] | Streamline lengths of 821 subjects from 2 cohorts averaged for 3 parcellations | 1 |
|  | Functional connectivity | [39] | Resting-state connectivity of 821 subjects from 2 cohorts | 5 |
|  | Functional connectivity | [37] | Anatomo-functional connectivity of 613 patients, 39 features | 1 |
| functional | BOLD time series | [24] | Streamline counts of 821 subjects averaged for 3 parcellations | 6 |

## Footnotes

1 Note that *siibra* also uses image-based assignment to brain regions for processing coordinates with significant localization uncertainty. This is realized automatically by converting uncertain coordinates into a 3D image of a Gaussian blob with corresponding kernel size.

2 The parcellation map may be hidden by clicking the “eye” icon near the parcellation selector, or the “y” key.

3 While not described in this article, it is possible to deploy the full tool suite including siibra-explorer and siibra-api on a local machine using siibra-compose (https://github.com/fzj-inm1-bda/siibra-compose). In this case, content added to the local configuration becomes immediately available in the local instance of siibra-explorer.

4 A command line client and Matlab^©^ library are being developed as well, but not covered in this contribution.

